# A bounded accumulation model of temporal generalization outperforms existing models and captures modality differences and learning effects

**DOI:** 10.1101/2024.10.15.616846

**Authors:** Nir Ofir, Ayelet N. Landau

**Affiliations:** Department of Psychology, Hebrew University of Jerusalem, Mt. Scopus, Jerusalem 9190501, Israel; Department of Cognitive and Brain Sciences, Hebrew University of Jerusalem, Mt. Scopus, Jerusalem 9190501, Israel; Edmond and Lily Safra Center for Brain Sciences, Hebrew University of Jerusalem, Edmond J. Safra Campus, Jerusalem 9190401, Israel

## Abstract

Multiple systems in the brain track the passage of time and can adapt their activity to temporal requirements (Paton & Buonomano, 2018). While the neural implementation of timing varies widely between neural substrates and behavioral tasks, at the algorithmic level many of these behaviors can be described as bounded accumulation (Balcı & Simen, 2024). So far, from the range of temporal psychophysical tasks, the bounded accumulation model has only been applied to temporal bisection, in which participants are requested to categorize an interval as “long” or “short” (Balcı & Simen, 2014; Ofir & Landau, 2022). In this work, we extend the model to fit performance in the temporal generalization task, in which participants are required to categorize an interval as being the same or different compared to a standard, or reference, duration (Wearden, 1992). Previous models of performance in this task focused on either the group level or performance of highly trained animals (Birngruber et al., 2014; Church & Gibbon, 1982; Wearden, 1992). Whether the same models can fit performance from a few hundreds of trials of single participants, necessary for comparing performance across experimental manipulations, has not been tested. A drift-diffusion model with two decision boundaries fits the data of single participants better than the previous models. We ran two experiments, one comparing performance between vision and audition and another examining the effect of learning. We found that decision boundaries can be modified independently: While the upper boundary was higher in vision compared to audition, the lower boundary decreased with learning in the task.

## Introduction

Accurately tracking the passage of time is crucial for all behavior. When we play a ball game, timing is critical from multiple perspectives. From a sensory perspective, we need to constantly track the ball and players and predict their next move. From a motor perspective, we need to time our hands and feet to meet the ball at the right moment. Neurobiological and theoretical studies suggest the implementation of tracking time takes many different forms, depending on the specific neural network and behavioral goal (Paton & Buonomano, 2018). Despite this variability in implementation, at the algorithmic level timing behaviors can often be described as a bounded accumulation process (Balci & Simen, 2016). The bounded accumulation model was first proposed for animal timing tasks, such as the peak interval procedure (Simen et al., 2011). In those tasks, an animal subject is given reward for the first response made after a certain interval has elapsed since a cue. In this case the model is most intuitive, as behavior can be explained by an accumulator and a decision boundary. The accumulator starts with cue presentation, and once it reaches the decision boundary, a response is made. The bounded accumulation model was later expanded to fit data from the temporal bisection task, a prototypical psychophysical timing task (Balcı & Simen, 2014). In this task, participants categorize intervals as being “short” or “long” based on two reference intervals they are familiarized with at the start of the experiment. With a minimal change, the bounded accumulation model can also fit data from this experiment: Instead of triggering a response, crossing the decision boundary signifies that an interval is long enough to be considered “long”. This model can explain both binary “short”/”long” responses as well as the EEG activity evoked by the offset of the interval (Ofir & Landau, 2022). Given the success of the bounded accumulation model in explaining behavior and EEG in temporal bisection, we wondered if it could accommodate other temporal tasks too. Specifically, can bounded accumulation explain behavior when more than one boundary is required? An example of such a task is the temporal generalization task, introduced to human research by John Wearden (1992). We first survey existing models, then describe an extension of the bounded accumulation model to this task. We next compare all models, showing the bounded accumulation model generally fits single participants’ data better than other models and has better parameter recovery. Finally, we demonstrate the use of the proposed model by investigating two scenarios. First, we use it to explain differences in temporal generalization between modalities. Second, we use it to account for changes in performance over learning.

## Method

### Participants

A total of 85 individuals participated in two experiments. Forty participated in experiment 1 (32 female participants, average age 23.9, SD 3) and 45 in experiment 2 (32 female participants, average age = 23.6, SD 2.6). Participants were recruited from the university community and were compensated for their time with either money (10 euro per hour) or class credit. All procedures were approved by the institutional review board of ethical conduct. Six participants from each experiment were excluded from the analysis (15% and 13.3%, respectively), as they produced flat psychometric curves, meaning they were not responding to the presented intervals (**Figure S1 & S2**).

### Experimental procedure

A typical temporal generalization experiment consists of several blocks of trials with an identical structure (**Figure 1**). Each block starts with several repetitions of the standard duration, which participants are not requested to respond to (Familiarization phase). Next, the test phase starts. In each trial of this phase, the participant is presented with a single interval and is asked to report whether the interval had the same duration as the standard or not. The presented duration varies from trial to trial, and usually covers a range spread evenly around the standard (**Figure 1B**).

**Figure 1:**
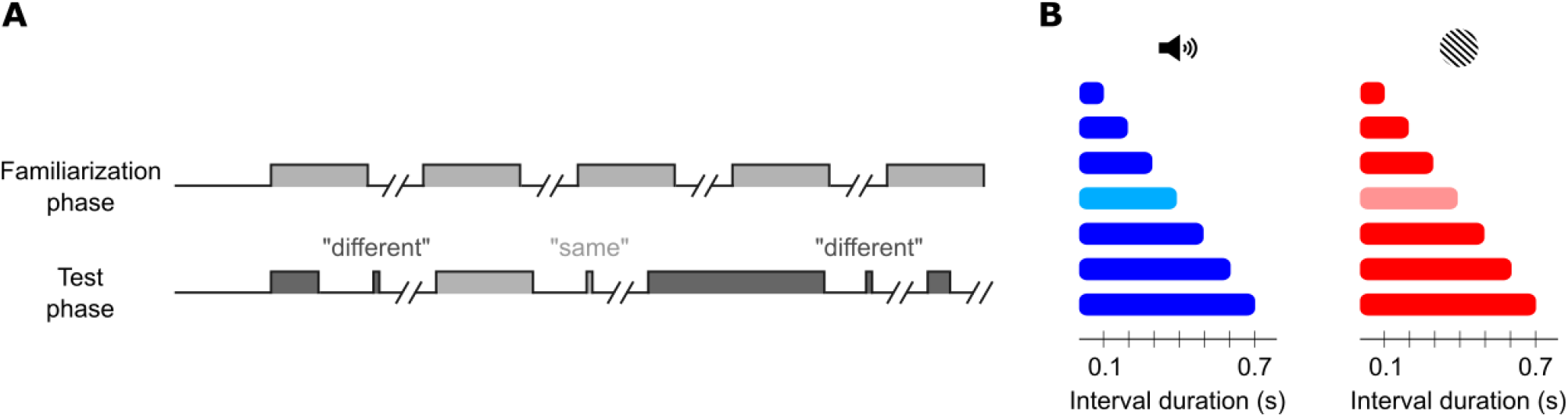
Schematic representation of a temporal generalization task. **A** Progression of the familiarization and test phases. While the familiarization phase is passive, after each interval in the test phase, the participant must respond before the experiment continues. **B** Intervals used in this work. In the first experiment, participants performed the same task twice, once with auditory stimuli and once with visual stimuli, in separate blocks. 0.4 s, in light color, is the standard.

We report the results of two behavioral experiments, both run using OpenSesame (Mathôt et al., 2012). The first compared temporal generalization with visual vs. auditory stimuli, and the second examined the effect of learning in the task. Both experiments used 400 msec as the standard duration, and seven levels of stimulus duration as comparison stimuli (100, 200, 300, 400, 500, 700 and 800 msec).

In the first experiment, participants completed two parts, one containing visual stimuli and one containing auditory stimuli. The order of the parts was counterbalanced across participants. Each part included 3 blocks of 75 trials each, separated by breaks. The standard duration was presented 5 times at the start of each block. At the end of a block, the percent of accurate responses was presented on the screen. In total, all durations except the standard were presented 30 times and the standard 45 times in each modality. A white fixation dot appeared at the center of the screen whenever no stimuli were presented on the screen.

In the second experiment we used visual stimuli in two levels of contrast (50% or 100%). The experiment contained 6 blocks of 80 trials each, separated by breaks. The standard duration was presented 6 times at the start of each block (3 in each contrast). At the end of a block, the percent of accurate responses was presented on the screen. All durations except the standard were presented 60 times (30 in each contrast) and the standard 120 times (60 in each contrast). Each block contained the same number of trials in each duration, displayed in a different order, to facilitate studying learning effects. A white fixation dot appeared at the center of the screen whenever no stimuli were presented on the screen.

In both experiments, participants could only respond once the stimulus was over.

### Experiment 1 stimuli

Visual stimuli consisted of a square-wave grating presented in a circular window on a BenQ XL2420Z monitor running on 144 Hz, which was positioned 50 cm away from the participants. The grating had a spatial frequency of 1 cycle per centimeter, a diameter of 7 centimeters (corresponding to 8° visual angle) and was positioned at the center of the screen. Gratings were presented with a random orientation of 45° or 135°. Auditory stimuli were 500 Hz tones presented at a comfortable hearing level via Sennheiser HD 280 pro headphones.

### Experiment 2 stimuli

Experiment 2 focused on the visual modality. The stimuli were the same square-wave gratings as presented in the visual part of experiment 1.

### Modelling temporal generalization performance

Developing mathematical models of behavior can serve several distinct goals (Wilson & Collins, 2019). One goal, which has guided early efforts to model temporal generalization, is to hypothesize and test the main sources of variability in performance (Church & Gibbon, 1982; Gibbon et al., 1984). This type of work sometimes places an emphasis on “well behaved” data, such as data pooled across participants, or the very large number of trials that come from experiments on animals. Another goal of computational modelling is to provide tools to summarize the data of single participants, which facilitates comparing behavior across conditions or individuals in cognitively meaningful terms (Schurr et al., 2024; Wilson & Collins, 2019). The second goal typically weighs more heavily the ability of the model to fit the “noisier” data of single participants.

All models considered here rely on a common approach in models of perception: a decision variable (DV) sample is drawn on each trial, and this sample is compared against two decision boundaries, to produce a binary decision: “Same” if the DV is within the boundaries, “different” otherwise. The models differ along several dimensions: Whether decision boundaries are constrained to be symmetric around the true standard duration, whether there’s trial-to-trial variability in the boundaries, and how noise in the internal representation of the current interval depends on the interval duration. In this work we focus on cognitive models for temporal generalization. Analysis approaches that assume a specific cognitive model were reviewed recently elsewhere (Bausenhart et al., 2018; See also Piras & Coull, 2011).

### Church & Gibbon 1982 (CG model)

Russel Church and John Gibbon originally developed a model to describe the performance of rats in a temporal generalization task (Church & Gibbon, 1982; Henceforth CG, following (Wearden, 1992)). The model assumes that on each trial a random sample of the standard duration is retrieved from memory as well as a random sample of the decision boundary. Then, the absolute value of the normalized difference of the current duration, which is assumed to be accurately perceived, and the sample of the standard is computed. This normalized difference is compared against the decision boundary. If the absolute normalized difference is smaller than the boundary, the interval is categorized as “same”, and if it is larger than the boundary, the interval is categorized as “different”. The decision rule can be formalized as follows:

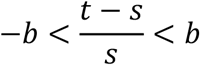

Where *s* (the standard memory) and *b* (the decision boundary) are both normally distributed random variables, and *t* is equal to the duration presented on the current trial. The psychophysical function – the probability to label an interval *t* as “same” – was derived by Church and Gibbon to be:

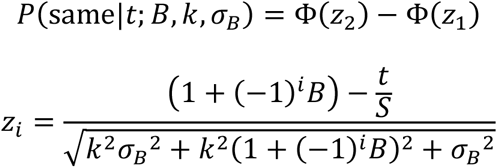

Φ is the standard normal cumulative density function and *S* is the true standard duration. The model has three free parameters: *B* (mean boundary value, relative to the standard), σ_*B*_ (boundary standard deviation) and *k* (Weber fraction, which specifies the variability of standard memory samples).

### Wearden 1992 (MCG: Modified Church & Gibbon model)

Later work that developed an analogue of the experimental design for humans found that humans displayed greater asymmetries in their psychophysical curves compared to rats (Wearden, 1992); Henceforth MCG “modified Church & Gibbon”). Wearden suggested to modify the DV, replacing the standard memoroy sample in the denominator with the objective duration of the current interval (the variables are the same as in the original model by Church & Gibbon):

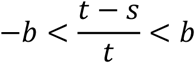

The psychophysical function can be found by algebraic operations (appendix 1) to be:

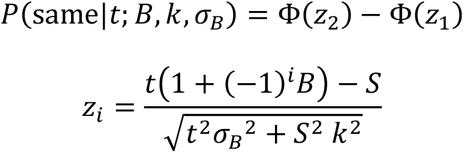

The free parameters are the same as for the CG model. Normalizing the DV by the interval duration instead of the standard means that the variability of the DV decreases for longer durations. This property is unusual for timing models, which typically assume that the uncertainty of estimated duration increases for longer intervals (Hass & Durstewitz, 2016). Over the years, many different variants of this model were developed to fit different scenarios, such as changes in performance over development (Droit-Volet et al., 2001; Wearden, 2004). We focus on the simplest one, as it provides the clearest comparison to the other models.

### Birngruber, Schröter & Ulrich 2014 (BSU model)

Both CG and MCG models assume that participants place their decision boundaries around the true standard duration. This assumption seems too strict for two reasons: First, individual participants often display idiosyncratic biases, in timing and other forms of perception, that result in shifted psychometric curves (Gibbon et al., 1984; Lebovich et al., 2019). Second, some experimental manipulations can create systematic shifts in the psychometric curves across participants. The third model we review was developed for data representing such a scenario. Birngruber and colleagues found that when the comparison interval is an oddball in a sequence of stimuli, it is perceived as longer than its objective duration (Birngruber et al., 2014; Henceforth BSU). Specifically, the peak of the psychometric function, which is the interval that is most often reported to be equal to the standard, is shorter than the standard. To allow the model to capture such biases, the authors suggested the following psychophysical function:

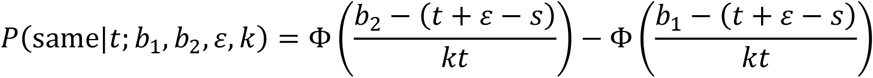

Where *s* and *t* are the objective standard and comparison intervals, respectively, *k* is the Weber fraction, which describes how quickly noise grows with the comparison interval, *b*_1_ and *b*_2_ are the lower and upper boundaries, respectively, and ε represents a bias term. We make two technical notes. First, as *s* is the objective standard duration, its only effect is that the boundaries are expressed as relative to the standard rather than in absolute terms. That can be done independently of the fitting procedure if desired. Second, ε trades-off perfectly with the boundaries. For any choice of ε, we can define new boundaries 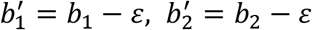 which would undo the effect of ε. In the original work, the value of the bias was constrained by additional data from a separate temporal bisection task. However, when only data from a temporal generalization task is available, this function is over-parameterized, and not all parameters can be estimated (see also appendix 2 in Birngruber et al., 2014). Hence, we remove ε and *s* from the function, yielding the simplified form:

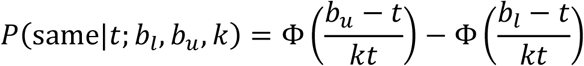

This model essentially states that a noisy estimate of the comparison interval is compared to 2 boundaries. If it is within those boundaries, it is reported as “same”, and as “different” otherwise. The model has three free parameters: The upper (*b*_*u*_) and lower (*b*_*l*_) boundaries and the Weber fraction *k*.

### Proposed drift-diffusion model (DDM)

Previous research showed that the bounded accumulation framework captures behavioral as wells as different aspects of neural activity in the temporal bisection task (Balcı & Simen, 2014; Ofir & Landau, 2022). We propose a modified drift-diffusion model (henceforth DDM), derived from this framework, for the temporal generalization task. The proposed model includes a single drift-diffusion process with two boundaries. Different from the typical implementation of the DDM in two-choice scenarios (Ratcliff et al., 2016), here both boundaries are placed above the starting point of the drift diffusion process. At interval onset, the drift-diffusion process starts, and the accumulated value is compared to the boundaries at interval offset. If the accumulated value at the offset hasn’t reached the lower boundary or has surpassed the upper boundary the interval is categorized as “different”. Otherwise, if the accumulated value at the offset is between the two boundaries, the interval is categorized as “same”. The psychophysical function is (see appendix 2 for the mathematical derivation):

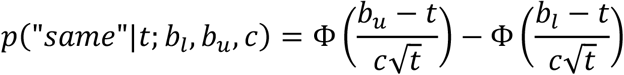

The model has three free parameters: the diffusion-to-drift ratio *c*, controlling how rapidly noise grows with time, the ratio of the lower boundary to drift (*b*_*l*_) and the ratio of the upper boundary to drift (*b*_*u*_). For brevity, the parameters will be denoted as diffusion coefficient, lower and upper boundary. We note that the DDM and BSU are very similar. They differ only in how rapidly timing noise grows with interval duration. The faster growth in variability assumed by BSU translates into curves that are generally more asymmetric than the ones produced by the DDM.

### Models summary

To summarize, all models assume a decision variable which is compared against decision boundaries. The DDM is the only model that explicitly describes the dynamics of the decision variable over time, so it is the simplest to plot as an example (**Figure 2A**). We can also think about the models through the psychometric curves they produce (**Figure 2B**). All four models have parameters that control the slopes (rising and falling) and asymmetry of the curve, which reflect the internal noise in the perceptual decision process. While the DDM and BSU models restrict noise to originate only from the timing process, CG and MCG assume that both timing (through the memory of the standard) and decision variability affect the slopes. In addition, DDM and BSU can produce curves that are not centered on the true standard, while CG and MCG cannot.

**Figure 2:**
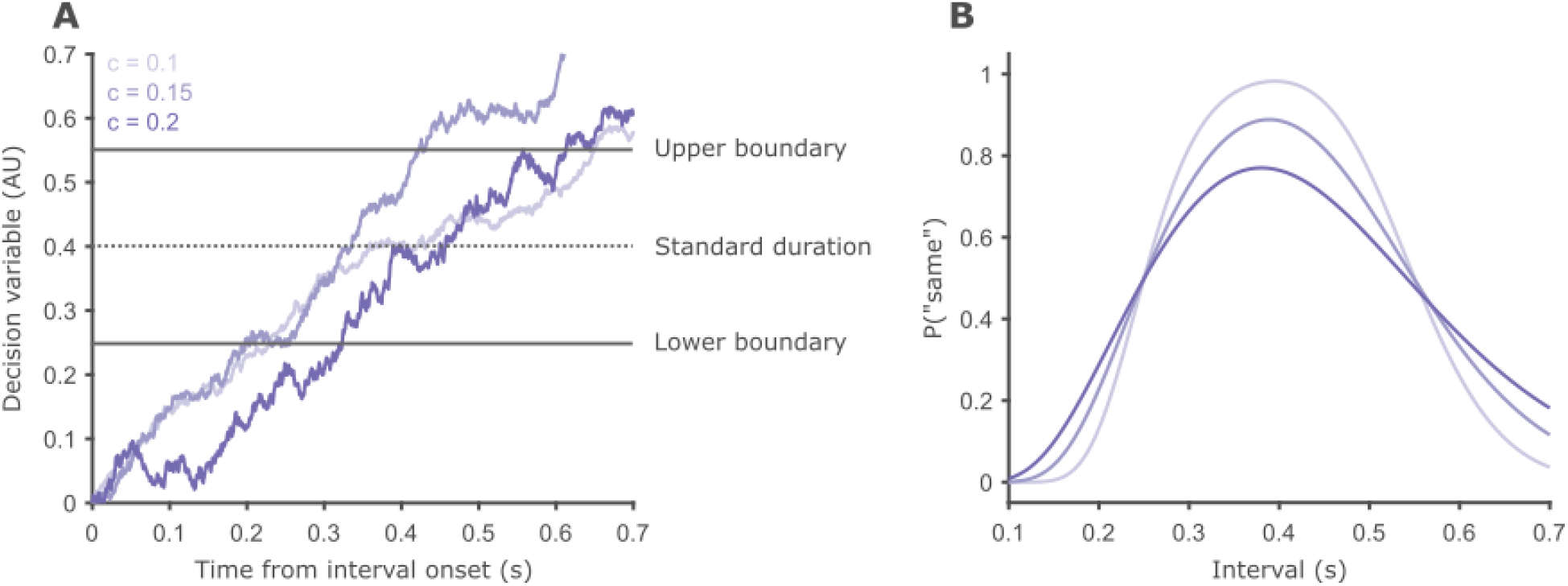
Schematic representation of the drift-diffusion temporal generalization model. **A** The basic components of the model – decision boundaries and decision variable – and examples of the evolution of the decision variable over time. Darker hues correspond to traces simulated with larger diffusion coefficients. Note the traces become more jagged as the diffusion coefficient increases. **B** Psychometric curves for the three levels of diffusion coefficient plotted in **A**. As internal noise grows, the curves become wider with shallower slopes, and the asymmetry increases.

### Fitting the model to behavior

The three free parameters of each model were estimated by a maximum likelihood procedure, using the fminsearch function of Matlab (MathWorks, MA). We used heuristics to make sure the fitting procedure started at parameter values that produce finite likelihoods (Wilson & Collins, 2019).

When working on the fits of the CG and MCG models, we noticed that fminsearch would sometime try combinations of the two slope parameters that lead to imaginary numbers in the denominator of the psychometric function, which cause Matlab errors. Therefore, for MCG and CG specifically, we constrained all parameters to be positive using fmincon.

### Parameter recovery

An important step in testing a model is examining its ability to fit data it simulated. This is called parameter recovery, and it measures the fitting capability of the model under an ideal situation (Wilson & Collins, 2019). In a parameter recovery analysis, a dataset is simulated by a model with given parameters, and then the model is fitted to the simulated data to estimate the model’s parameters. If the model parameters are well defined, and the data collection is well suited, we expect that the estimated parameter values will be close to the values used to simulate the data. We created parameter-generating distributions for the simulations by fitting a probability distribution to the fitted values of each parameter separately (twelve independent distributions in total, three for each of the four models). Fits of the MCG and CG models had four outlier datasets each, that were removed prior to the parameter recovery analysis. We chose the distributions manually to reasonably fit the parameter values. For each model, 5000 simulations were generated from the parameter distributions using the same trial numbers as in the actual experiment. The simulated parameters were independent, except for the upper and lower boundaries in the DDM and BSU. Simulations in which the upper boundary was less than 0.1 larger than the lower boundary were redrawn. Finally, the results of each simulation were fitted by the model that created it, and the estimated and simulated parameters were compared (**Figure 6**). As a general measure for fitting accuracy, we calculated the Pearson correlation coefficient between simulated and estimated parameter values. Parameter trade-off was assessed using the correlation between all pairs of estimated parameter values.

## Results

### The double-boundary DDM fits single participants’ data better than the other models

We analyzed the data of 34 participants who completed two versions of temporal generalization, one block using auditory pure tones and one using visual gratings. Overall, participants were better with auditory stimuli (**Figure 3**). Participants performed significantly more accurately in the auditory modality (M = 71.01%, SD = 9.52%) than in the visual modality (M = 58.77%, SD = 8.88%). All participants but one had higher accuracy in the auditory modality (paired t-test, t_33_ = 11.18, p < 0.001, d = 1.92). We performed a repeated-measures analysis of variance (ANOVA) on the probability of “same” responses with interval duration (7 levels, 0.1-0.7 s), modality (2 levels, audition or vision) and their interaction as within-participant factors. We found an expected significant main effect of interval duration 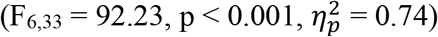, signifying that participants were attending to the task. In addition, we found a significant main effect of modality 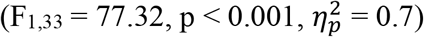, as well as a significant duration by modality interaction 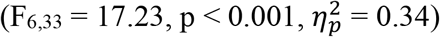. Visually inspecting participants’ performance shows that the psychometric curves for auditory and visual stimuli differ greatly in their shapes. These differences between are most pronounced for longer intervals. Considering Weber’s law, this would intuitively correspond to a larger coefficient of variation in vision compared to audition: Timing noise for static gratings grows at a faster rate compared to pure tones. To test that intuition formally and statistically, we can use cognitive models, as explored below.

**Figure 3:**
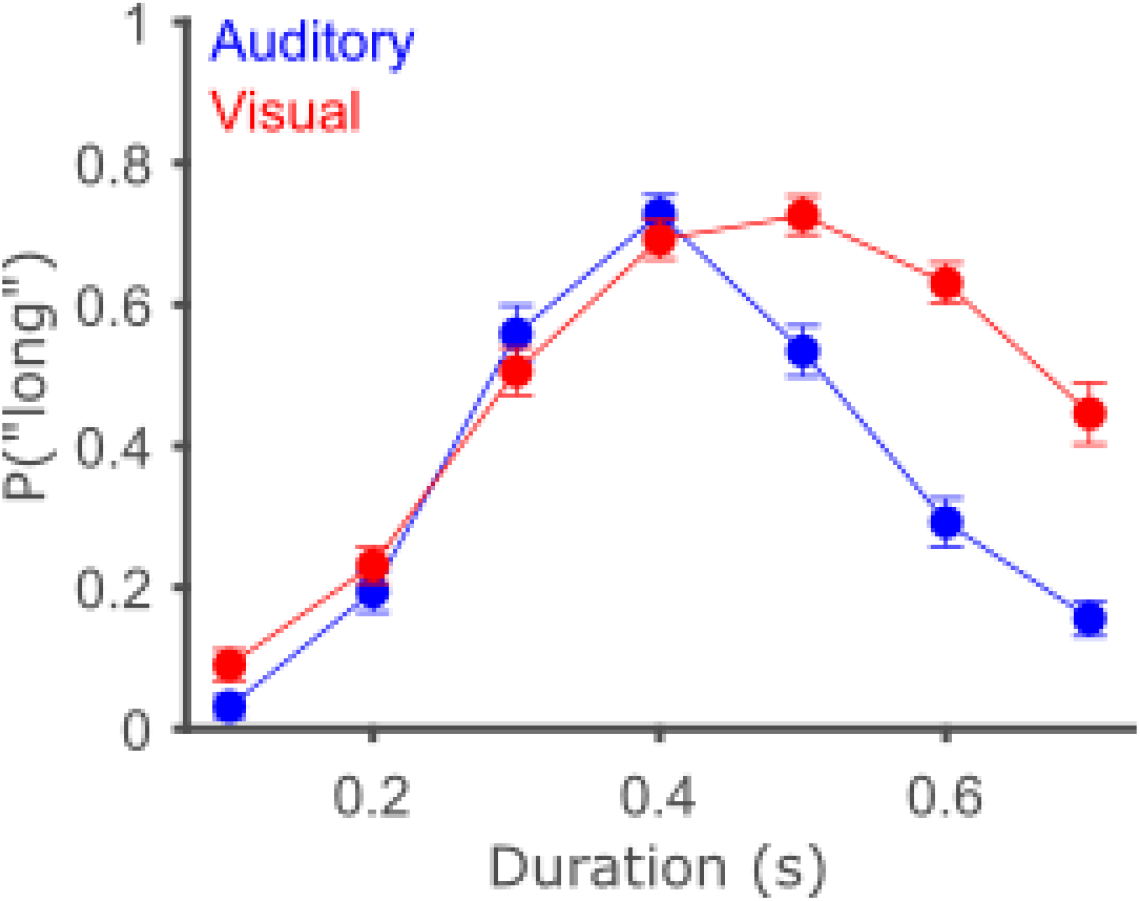
Participants perform better on auditory intervals. Circles show the average probability to label an interval as “same*”* across participants, and error bars depict the within-participant SEM using the Morey-Cousineau method (Cousineau et al., 2021). In blue is data from the auditory modality and in red data from the visual modality.

Comparing behavior under different conditions is often done by first summarizing behavior within conditions into model parameters, and then comparing the parameter values between conditions. To do so, we need to establish the models are suitable, both in how well they can fit the data and in how well-defined the models’ parameters are. First, we compared the ability of the four different models to fit data at a single participant level. For each participant, we fit each model to the data of each condition separately. Since all models have three free parameters, they can be directly compared in terms of their maximum likelihood. For both modalities, the DDM considerably outperformed all other models (**Figure 4**), with 47.1% and 73.5% of participants in the auditory and visual modalities respectively, compared with a chance level of 25% (**Table 1**). There was no obvious systematic difference between the other three models. Models BSU and CG were somewhat better on auditory than visual data. MCG performed the same in both modalities. Following previous research, we also fitted the models to the pooled data across participants. In the auditory modality, the original CG model performed best, while in the visual modality it was the MCG model. In both modalities, the DDM came in second. Visually inspecting the group results reveals the CG model tracks the auditory data closely, while the MCG model tracks the visual data closely (**Figure 5**). Both models failed quite clearly in fitting the other modality (i.e. CG in the visual modality, and MCG in the auditory modality). The DDM was more balanced in fitting the group data, achieving good yet not optimal fits in both modalities. Importantly, the fits at the group level should be taken with a grain of salt. Accepting the group results at face value assumes that the pooled data is a cleaner version of the prototypical pattern of performance, or in other words, that participants behave similarly. However, participants in psychophysical tasks often display significant variability (**Figure S1**; See also Lebovich et al., 2019). In more mathematical terms, given the variability of single participants, there is no guaranty that the same psychometric function applies at both single participant and group levels. As reported above, at the single participant level, the DDM considerably outperforms the other models.

**Table 1:** Number of participants and group log-likelihood for each model in each modality. Bold numbers mark the model that performed best in each modality at the single participant or group level.

|  | Audition |  |  | Vision |  |  |
| --- | --- | --- | --- | --- | --- | --- |
|  | Nr. participants | % of participants | Group LL | Nr. participants | % of participants | Group LL |
| DDM | **16** | **47.1** | -4045.9 | **24** | **70.6** | -4530.7 |
| BSU | 7 | 20.6 | -4221.5 | 4 | 11.8 | -4607.9 |
| MCG | 4 | 11.8 | -4183.2 | 4 | 11.8 | **-4513.8** |
| CG | 7 | 20.6 | **-4008.9** | 2 | 5.9 | -4618.5 |

**Figure 4:**
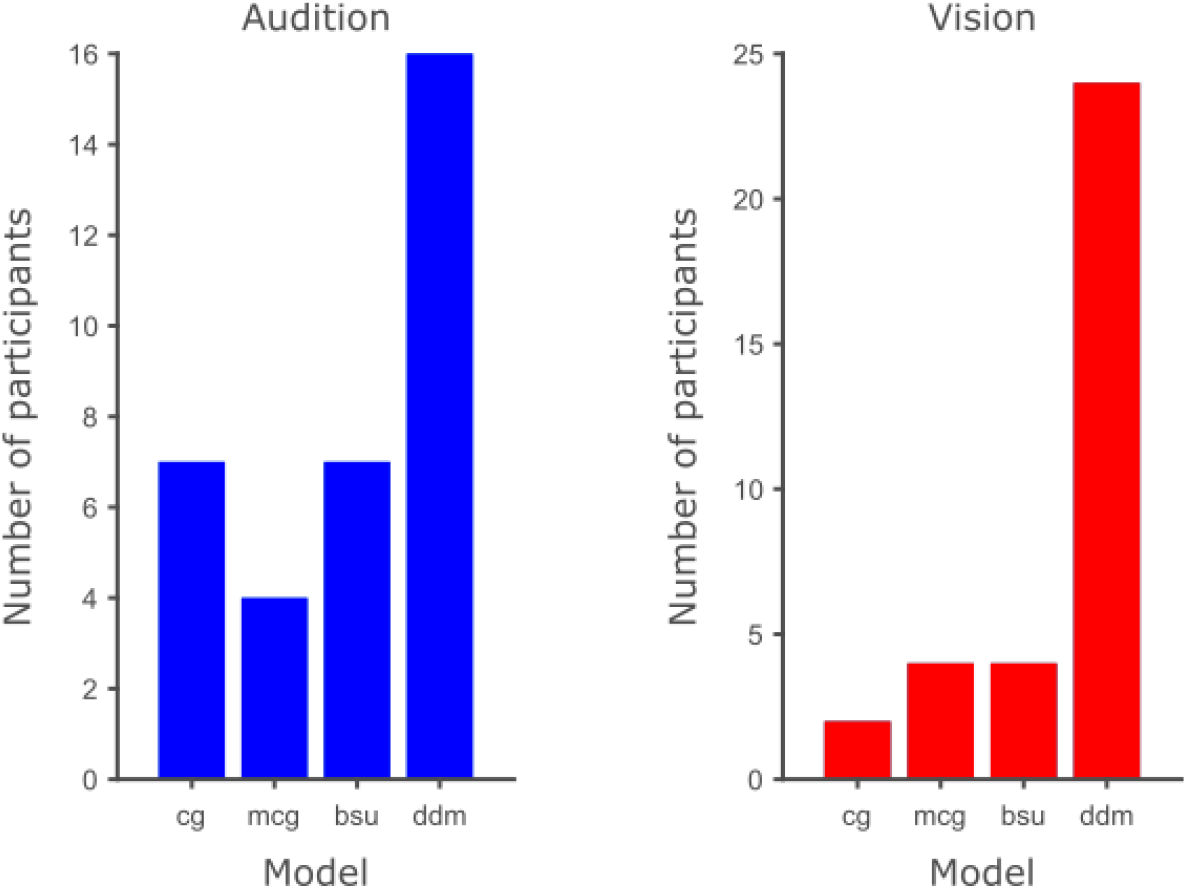
The DDM outperforms the other models at the single participant level in both modalities. The height of each bar signifies the number of participants (out of a total of 34) for which each model, at the x-axis, achieved the largest likelihood. Left shows results from the auditory block and right shows data from the visual block.

**Figure 5:**
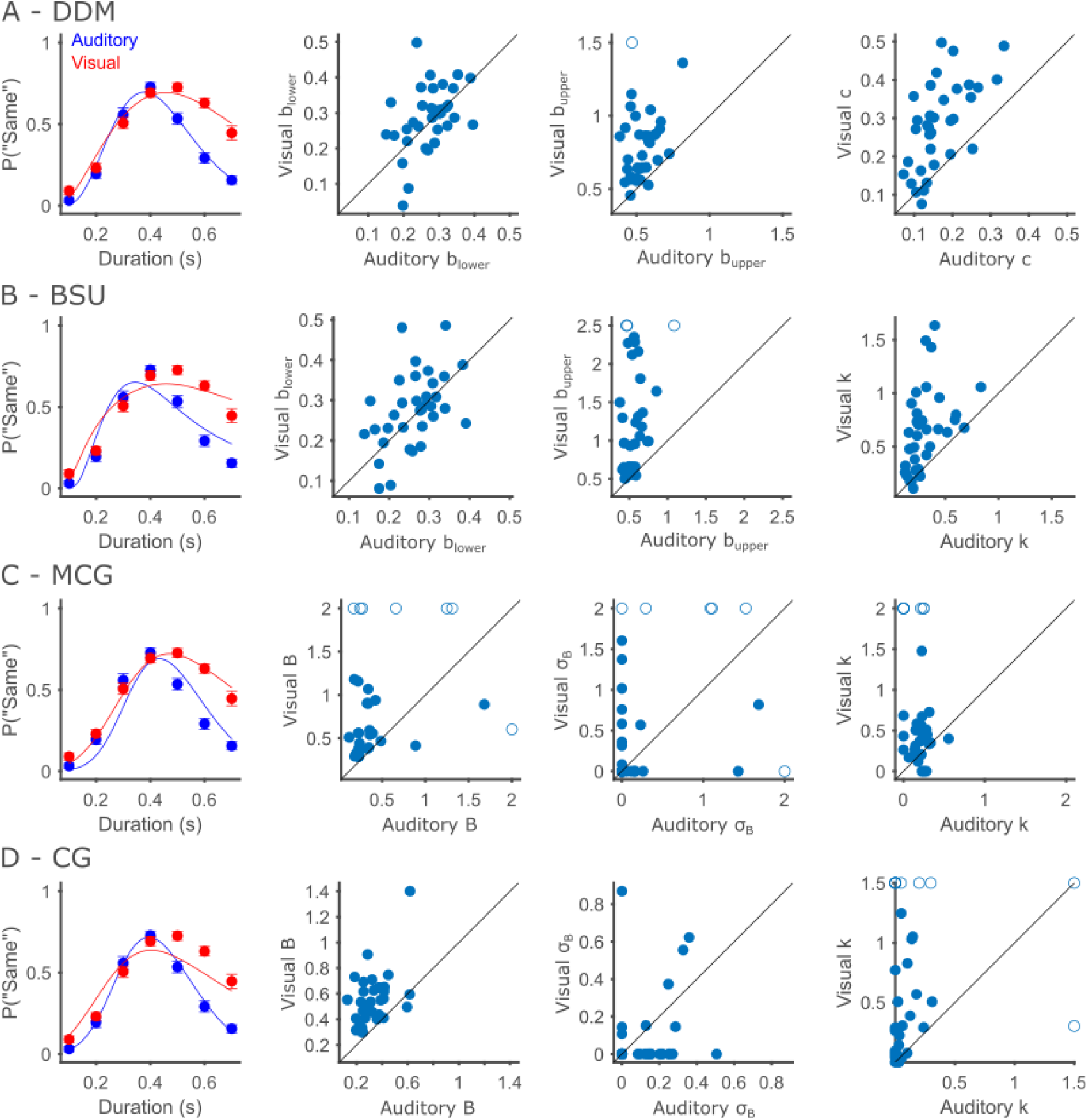
Behavioral performance and model fits for Experiment 1. Each row shows results of one model. **A** DDM. Leftmost is the performance and model fits at the group level. Circles show the average probability to label an interval as “same*”* across participants, and error bars depict the within-participant SEM. Scatter plots show the estimated values of each of the free parameters (Left to right: lower boundary, upper boundary and diffusion). X and Y axes correspond to the auditory and visual modalities, respectively. To facilitate visualization, parameters far from the group are truncated and depicted with empty circles. **B** Same as **A**, for the BSU model. Parameter scatter plots show, left to right: lower boundary, upper boundary and Weber fraction. **C** Same as **A**, for the MCG model. Parameter scatter plots show, left to right: Mean boundary, boundary standard deviation and Weber fraction. **D** Same as **A**, for the CG model. Parameter scatter plots show, left to right: Mean boundary, boundary standard deviation and Weber fraction.

**Figure 6:**
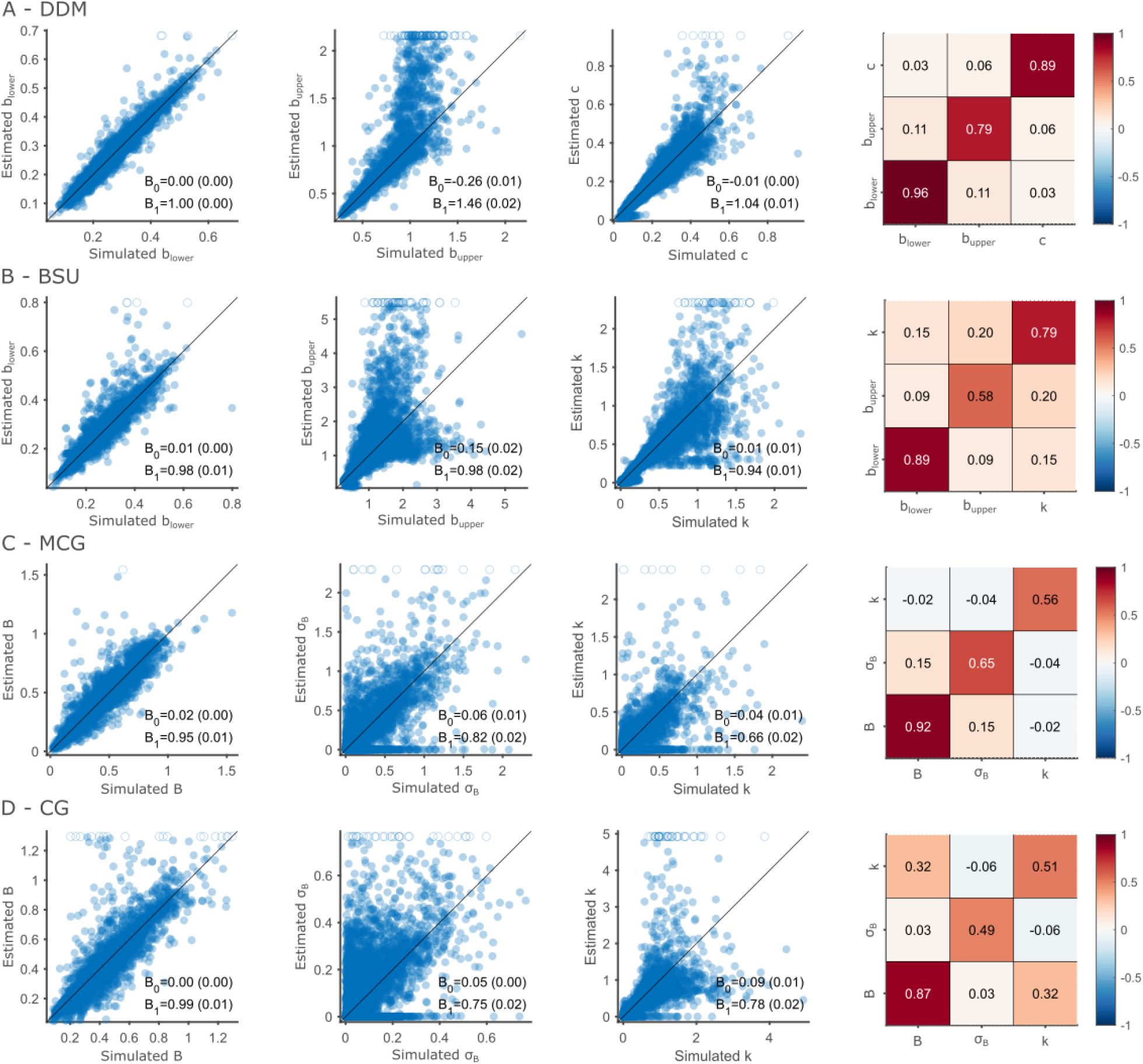
DDM Parameters are more accurately recovered than the other models. Each row shows the result of one model. **A** Parameter recovery for the DDM. Scatter plots show the estimated parameter (y-axis) against the simulated parameter (x-axis), for each of the free parameters (left to right: lower boundary, upper boundary and diffusion). Each point represents a single simulation. To facilitate visualization, values that were estimated as larger than the largest simulated parameter are truncated and replaced with empty circles. The heatmap at the right show the parameter correlations. Diagonal cells depict the correlation coefficient between simulated and estimated parameters, and off-diagonal cells depict the correlation between the estimated values of two different parameters. **B** Same as **A**, for the BSU model. Scatter plots depict, left to right: lower boundary, upper boundary and Weber fraction. **C** Same as **A**, for the MCG model. Parameter scatter plots show, left to right: Mean boundary, boundary standard deviation and Weber fraction. **D** Same as **A**, for the CG model. Parameter scatter plots show, left to right: Mean boundary, boundary standard deviation and Weber fraction.

As noted, both CG and MCG models have two parameters that control the slopes and asymmetry of the curves: Boundary variability and memory variability, controlled by the Weber fraction. Having more than one parameter affecting the psychometric curve similarly could lead to identifiability problems, where changes to one parameter can be undone by changes to another parameter (Gershman, 2016). This means both parameters cannot be reliably estimated from typical empirical data at once. Identifiability is especially important if the parameters are used for inference, such as comparing between experimental conditions. To test parameter identifiability, we explored the estimated parameter values of all models (**Figure 4**). Fits of both CG and MCG models show a tendency to shrink one of the two parameters towards zero, more often boundary variability, leaving the other parameter to absorb all explained variability (**Table 2**). This suggests the slopes parameters are unidentifiable. We note this was already reported briefly by Wearden (1992). The CG model was only used to fit pooled data of several animals, each completing many hundreds of trials. These large amounts of data, atypical in human psychophysics, possibly allowed the fitting procedure to distinguish both sources of variability (Gibbon et al., 1984). However, for the type of data discussed here, these models are suboptimal.

**Table 2:** Parameter unidentifiability in the MCG and CG models. Each cell includes the number of participants (out of 34) for which the specific parameter (σ_*B*_, *k* or both) was estimated to be smaller than 0.001.

|  | MCG |  |  | CG |  |  |
| --- | --- | --- | --- | --- | --- | --- |
| | $\sigma_B$ | $k$ | Both | $\sigma_B$ | $k$ | Both |
| Audition | 21 | 6 | 0 | 15 | 12 | 0 |
| Vision | 19 | 3 | 0 | 26 | 4 | 0 |

### Parameters of the DDM are more successfully recovered than the other models

Finally, we tested the validity of the fitted parameter values as estimates of the values which hypothetically produced the data by parameter recovery (Wilson & Collins, 2019). The DDM model displayed the overall highest recovery accuracy for all three parameters. Upper boundaries were generally estimated well, up to upper boundaries of about 0.8 s. Above this point, the fitting procedure tended to inflate the estimated upper boundaries. This means upper boundaries estimates larger than 0.8 should be treated somewhat cautiously. In our data, estimated upper boundaries above 0.8 s occurred only once in the auditory condition, but 17 times (50% of participants) in the visual condition. This reveals a limitation in the experimental design. It is possible that the range of intervals used in the experiment was too difficult for many of our participants in the visual modality. Cross-parameter correlations were low overall, indicating good parameter identifiability, except for the correlation between lower and upper boundaries. As the simulated boundaries were constrained to have a separation of at least 100 ms, there was a correlation of 0.15 in the simulated values, close to the 0.13 correlation in the estimated values. The BSU model displayed somewhat worse parameter recovery, which is most apparent in the estimation of upper boundaries in that model. The variability of estimated upper boundaries increased quite rapidly with increasing simulated upper boundaries. This translates into lower recoverability, as measured by the correlation between simulated and estimated upper boundaries. For both MCG and CG models, mean boundary was recovered well. However, both slope parameters showed low recovery accuracy across the entire range of values tested. The trade-off between slope parameters, which was very strong when fitting empirical data, was less severe in the recovery analysis. This is evident in the wide spread of estimated slope values. This suggests that the trade-off found in fitting single participants data stems from the models being generally less suitable for such data. In summary, the DDM showed satisfactory parameter recovery, better than the other models explored, as well as minimal trade-off between parameters, making it a good candidate to test the effect of experimental manipulations.

### Participants have less internal noise and use stricter decision boundaries when timing auditory vs. visual stimuli

Having established the DDM as a suitable model to analyze single-participant data, we next use it to capture differences between conditions. We fit the DDM to the data of each participant within each condition separately (**Figure 5**) and compared the estimated parameters between the conditions using paired t-tests. Lower boundaries weren’t significantly different between modalities (M_aud_ = 0.27, SD_aud_ = 0.06, M_vis_ = 0.29, SD_vis_ = 0.09; t_33_ = 0.99, p = 0.329, d = 0.17). In contrast, both the diffusion and upper boundaries were strongly affected by the modality. Diffusion coefficients were significantly larger in the visual condition (M_aud_ = 0.17, SD_aud_ = 0.06, M_vis_ = 0.28, SD_vis_ = 0.11; t_33_ = 7.50, p < 0.001, d = 0.77), and participants placed their upper boundaries at longer intervals in the visual condition (M_aud_ = 0.54, SD_aud_ = 0.09, M_vis_ = 0.83, SD_vis_ = 0.38; t_33_ = -4.51, p < 0.001).

As both internal noise and upper boundary were significantly different between modalities, we next explored whether both parameters might be inherently related. If this were the case, both parameters should be correlated across participants. To make sure our statistical analysis isn’t overly affected by outlier values, we checked for upper boundaries or diffusion coefficients that were 3.5 standard deviations or more away from the respective means. We excluded 1 participant with an upper boundary of 2.6, which is 6.27 standard deviations above the mean upper boundary. We ran a linear mixed effects model predicting the upper boundary using modality (binary predictor with effects coding: -1 for audition and 1 for vision) and diffusion (continuous predictor) as fixed effects. The intercept and slope against diffusion were set as random effects. We found that participants with larger diffusion tended to place their upper boundary at longer intervals (β = 0.91, p < 0.001; **Figure 7A**). Modality still predicted significant variability in upper boundaries, even after taking diffusion into account (β = -0.07, p < 0.001). Additionally, the relation between diffusion and upper boundary was stronger in the visual modality, as indicated by a significant interaction of modality and diffusion (β = -0.38, p = 0.011). Importantly, the correlations seen in the empirical data are larger than those seen in the parameter recovery analysis, meaning this correlation reveals a true link between the two measures, rather than an artifact of the fitting procedure or model specification. If diffusion and upper boundaries are related, another prediction is that participants who displayed a larger effect of modality on their diffusion coefficient, should also display a larger effect of modality on upper boundaries. Hence, we computed the difference of diffusion coefficient between modalities (vision minus audition, so we generally expect positive differences) as well as the difference of upper boundaries and correlated the two. The two differences were significantly correlated (Pearson ρ = 0.56, p < 0.001; Figure 7B) in line with the prediction. In summary, upper boundaries and diffusion are related, yet the effect of modality on upper boundaries is not completely explained by its effect on internal noise.

**Figure 7:**
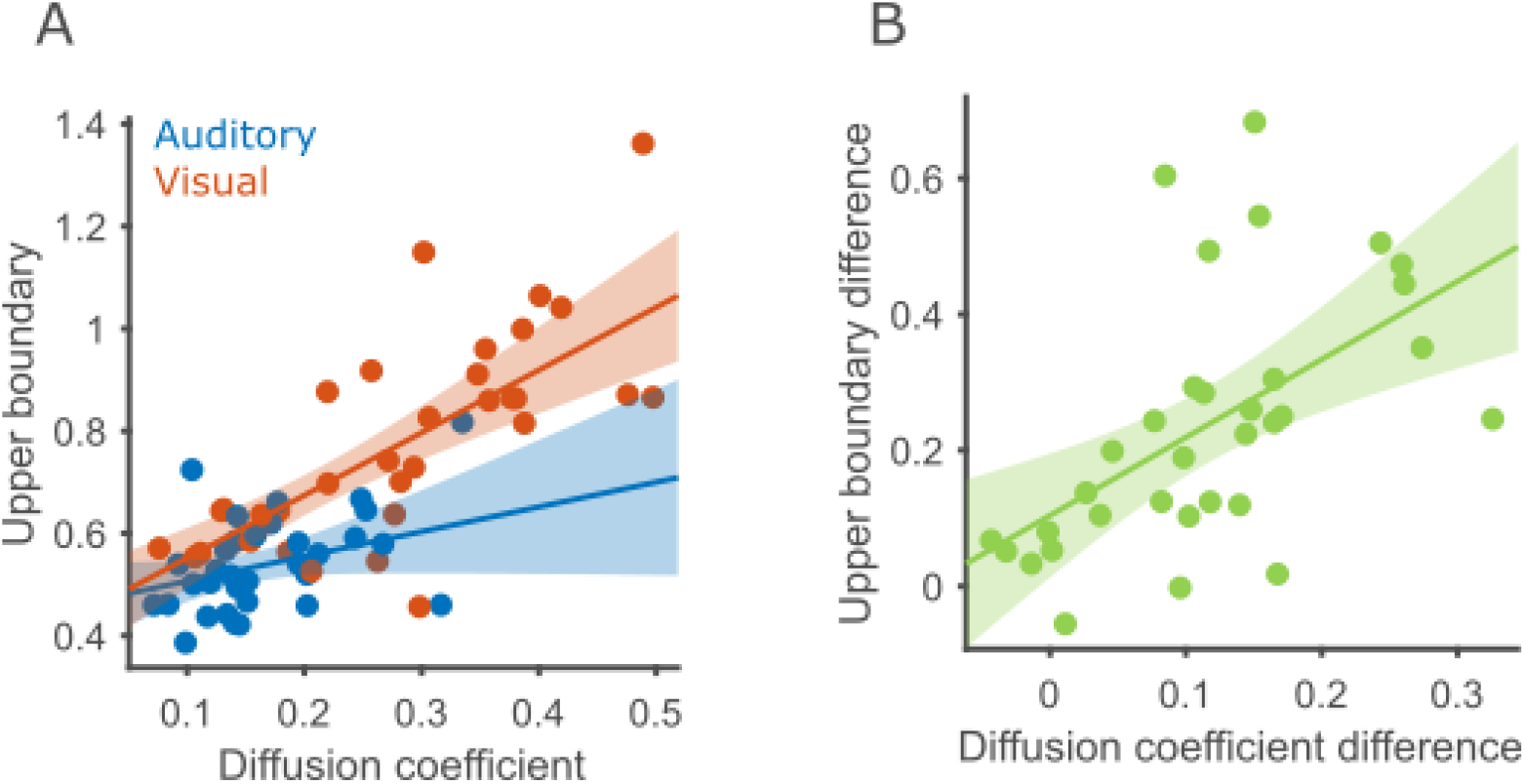
Diffusion coefficient and upper boundary are correlated. **A** Correlation of diffusion coefficient and upper boundary in both modalities. Circles represent the data of a single participant in a single modality (blue for auditory and red for visual). The 95% confidence interval of the regression lines are marked using shaded ribbons. **B** correlation across participants between upper boundary difference (vision – audition) and diffusion coefficient difference.

### Psychophysical functions become narrower with increased experience in the task

We analyzed the data of 39 participants in the second experiment, exploring the effect of learning in the task. We performed a repeated-measures ANOVA on the probability of “same” responses with interval duration (7 levels, 0.1-0.7 s), block (3 levels, blocks 1-2, 3-4 and 5-6) and their interaction as within-participant factors. As expected, there was a significant main effect of interval duration 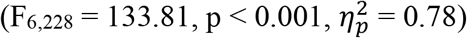. More importantly, we found a significant main effect of block number 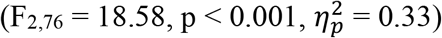, as well as a significant duration by block interaction 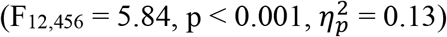. This means participants systematically modified their behavior over the course of the experiment. Visually inspecting participants’ performance shows that the generalization gradients became narrower with longer experience with the task and increasing exposure to the standard durations (**Figure 8**). We next examined whether participants adapted their decision strategy and whether their internal timing noise changed as a function of learning, by fitting the DDM to the data of each participant and block independently (meaning 39 × 3 models were fitted in total). We then performed a repeated-measures ANOVA on the estimated parameters with block (3 levels, blocks 1-2, 3-4 and 5-6) as a within-participant factor. We found that the lower boundary differed significantly between blocks 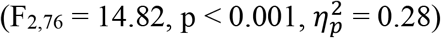, shifting to shorter durations with learning (Tukey-Kramer post-hoc tests, 1^st^ vs. 2^nd^ tertile: p = 0.010, 1^st^ vs. 3^rd^: p < 0.001, 2^nd^ vs. 3^rd^: p = 0.021). Diffusion coefficients also changed significantly between blocks 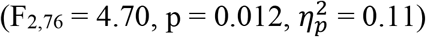, becoming somewhat smaller over the course of the experiment. A post-hoc test found a significant difference between the 1^st^ and 3^rd^ tertiles (1^st^ vs. 2^nd^ tertile: p = 0.275, 1^st^ vs. 3^rd^: p = 0.017, 2^nd^ vs. 3^rd^: p = 0.237). Upper boundaries did not significantly differ between blocks 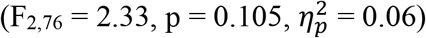.

**Figure 8:**
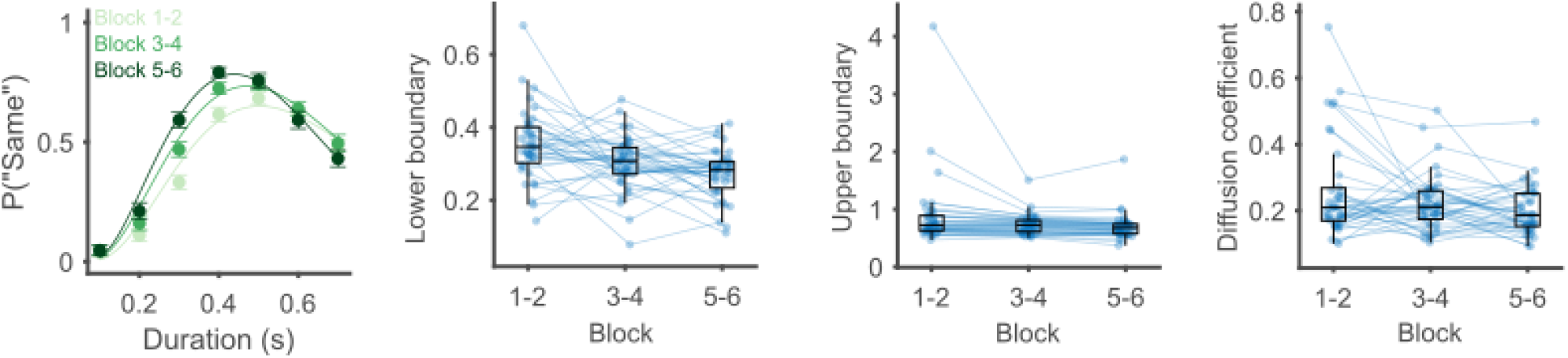
Psychophysical performance improves with learning. Group fits and individual parameter estimates for each block using the DDM. Left, performance and model fits at the group level. Circles show the average probability to label an interval as “same*”* across participants, and error bars depict the within-participant SEM. Scatter plots show the estimated values of each of the free parameters for data of each block. Circles and connecting lines mark the parameter values across blocks for a single participant, with box plots overlaid. Horizontal lines within the boxes represent the group median, the box extends from the 25th percentile to the 75th percentile, and the whiskers extend 1.5 IQRs from the median.

Here, as in the comparison between modalities, we found two parameters of the model - diffusion coefficient and lower boundary - that changed significantly between blocks. As before, we first tested for outlier values in lower boundaries and diffusion coefficients, with 3.5 standard deviations away from the mean as the rejection criterion. We removed one participant which had a diffusion coefficient in the first block 4.8 standard deviations above the mean. We next used a linear mixed effects model to predict the lower boundary using block (categorical predictor with the first block as the reference level), diffusion coefficient (continuous predictor) and the diffusion coefficient by block interaction as fixed effects and a random intercept per participant. As expected, lower boundaries were gradually shifted to shorter intervals over blocks (2^nd^ vs. 1^st^ tertile, β = -0.04, p = 0.003; 3^rd^ vs. 1^st^ tertile, β = -0.06, p < 0.001). The diffusion coefficient wasn’t significantly correlated with lower boundaries in the first tertile (β = 0.12, p = 0.165). The interaction terms were not significant either, suggesting similar relations of diffusion to lower boundaries over learning (2^nd^ vs. 1^st^ tertile, β = -0.02, p = 0.864; 3^rd^ vs. 1^st^ tertile, β = 0.16, p = 0.283; **Figure 9**). The mixed model results suggest that learning affects the lower boundary and diffusion coefficient independently.

**Figure 9:**
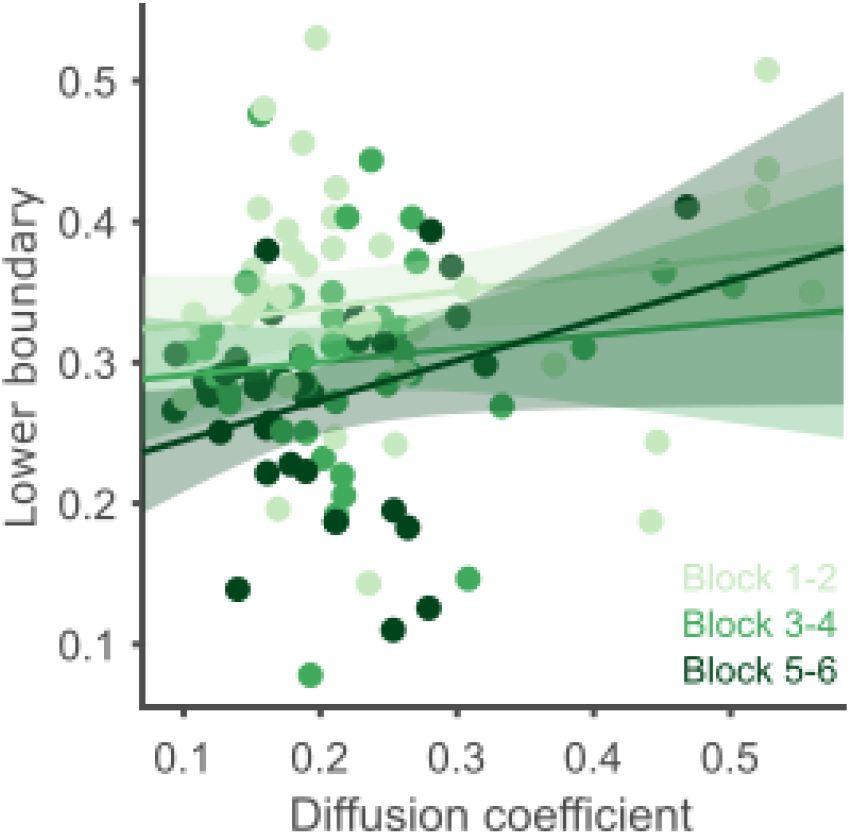
Diffusion coefficients and lower boundaries aren’t correlated. Correlation of diffusion coefficient and lower boundary in the three tertiles. Each circle represents the data of a single participant in a single tertile (Darker colors for later tertiles). The 95% confidence interval of the regression lines are marked using shaded ribbons.

## Discussion

A diverse set of neural mechanisms is thought to underly our capability to track the passage of time (Paton & Buonomano, 2018). Yet, at the algorithmic level many timing behaviors can be described as bounded accumulation processes (Balcı & Simen, 2024). The bounded accumulation framework has been successfully applied to behavioral performance in temporal reproduction tasks (Simen et al., 2011). An extension of the framework to more complex temporal decisions, such as temporal bisection, explains results at the level of behavior (Balcı & Simen, 2014) and EEG (Ofir & Landau, 2022). In temporal bisection, binary responses (“long” or “short”) and EEG patterns could be captured with a single decision boundary (Ofir & Landau, 2022). In this work, we extend the framework further to temporal generalization.

The previous models, as well as the proposed DDM, all share the basic design of psychophysical models, that is comparing a decision variable against decision boundaries, but vary considerably in their assumptions regarding the boundaries and the sources of variability. The original models, CG and MCG (Church & Gibbon, 1982; Wearden, 1992), constrained decision boundaries to be fixed around the true standard duration and assumed that performance variability in the task (i.e. the slopes of psychophysical functions) is driven by noise in the memory of the standard and the decision boundaries. A more recent model, BSU (Birngruber et al., 2014), relaxed the constraint on the decision boundaries, and removed their trial-to-trial variability. Here, we apply the drift-diffusion approach to temporal generalization. We defined two decision boundaries, free to take any values, as well as a diffusion coefficient controlling the level of internal noise.

The DDM outperforms all other models in fitting single participants’ data. This is found both for timing using auditory stimuli and using visual stimuli. Furthermore, an analysis of parameter recovery revealed that the DDM parameters were best recovered. The two slope parameters of the CG and MCG models were recovered especially inaccurately, meaning the values estimated from empirical data are less reliable for these two models.

Having a model which can be robustly fitted to single participants’ data enables statistically testing how experimental manipulations affect the cognitive processes underlying behavior. Using the proposed DDM, we found that timing visual stimuli differed in two ways when compared to timing auditory stimuli. First, participants had a noisier representation of time in the visual modality, reflected by higher diffusion coefficients. Second, they also adopted higher upper boundaries in the visual modality, meaning that they were more likely to categorize longer intervals as identical to the standard. Numerous previous studies found timing of auditory stimuli to be better than timing of visual stimuli, leading to the hypothesis that audition is more temporally accurate (Di Luca & Rhodes, 2016; Espinoza-Monroy & De Lafuente, 2021; Wearden et al., 1998). A post-hoc analysis revealed that participants with more internal noise placed their upper boundaries at longer intervals. This result is reminiscent of previous findings, which indicate that participants adapt their behavior according to their level of internal noise (Freestone & Church, 2016; Kononowicz et al., 2022). However, even when this correlation is considered, modality has an additional independent effect on upper boundaries, with upper boundaries placed at longer intervals for visual stimuli. These results suggest that modality affects timing at the perceptual level, with visual stimuli less precisely timed, as well as at the decision-making level, with less strict boundary placement for visual stimuli.

We additionally used the DDM to uncover the cognitive processes affected by learning in the task, using data from a second temporal generalization experiment. At the raw behavioral level, we found that psychophysical curves became narrower with increased experience with the task. This sharpening was captured in the DDM as a lowering of the lower decision boundaries, accompanied by a decrease in the diffusion coefficient. We found that participants initially placed the lower boundary almost at the true standard duration. As they gained practice with the task, participants gradually shifted their lower boundaries to shorter durations, which might reflect a more accurate representation of the standard. Our findings contrast with a previous study which suggested that upper boundaries were lowered with learning (Wearden & Towse, 1994). However, that study, which relied on the MCG model and a variant of it, did not statistically compare parameter estimates between blocks of trials. Additionally, their pooled behavioral results (Wearden & Towse, 1994; Figure 2) are much sharper than the ones we found, making direct comparisons difficult.

The results of the modality and learning experiments combined show that the lower and upper boundaries can be modified independently of each other. This contrasts with the symmetric boundaries assumed by the CG and MCG models, which implies both should change in concert. Whether the boundaries are symmetric or not, all four models assume they are stable across trials, or vary randomly from trial to trial. Past studies, about time perception as well as many other forms of perception, suggest decision boundaries and internal references change systematically across trials (Hachen et al., 2021; Raviv et al., 2012; Wiener & Thompson, 2015). Future work could leverage serial dependence to explore how the recent history of presented intervals and responses affects temporal generalization performance.

Finally, in the proposed model we assumed that the decision variable is compared against the boundaries only at interval offset, regardless of whether it crossed the boundaries during the interval. In the context of drift-diffusion models, such boundaries are termed “unabsorbing”. However, it is reasonable to hypothesize that the upper boundary is absorbing. In other words, relatively long intervals could be categorized as “different” before their offset, as soon as the decision variable reaches the upper boundary. An analogous process is thought to occur in temporal bisection, where “long” decisions are often made before interval offset (Ofir & Landau, 2022). Two predictions can be made based on this hypothesis, at the behavioral and physiological level. First, the fact that for longer intervals “different” decisions can be made before interval offsets predicts faster responses in those cases, as motor preparation can start in advance. There is some evidence that this is the case (Klapproth, 2018; Klapproth & Müller, 2008; Klapproth & Wearden, 2011). Second, breaking the decision process into two stages, until interval offset and after it, suggests that we can expect to see a similar pattern of neural signatures in temporal generalization and bisection. Specifically, it is expected that the offset-evoked potential in temporal generalization will be larger for shorter intervals, as it is in bisection (Ofir & Landau, 2022). To our knowledge, only two studies examined the offset-evoked potential in temporal generalization (Bannier et al., 2019; Özoğlu & Thomaschke, 2023). Both studies found the same pattern of non-linear decreasing EEG potential as a function of interval duration in temporal generalization and bisection. A DDM with absorbing upper boundaries would be a beneficial tool for future studies recording response times alongside non-invasive physiology to study the temporal dynamics of the decisions in the temporal generalization task.

In summary, we provide a model for temporal generalization, which can robustly fit single participants data, enabling us to directly test the effect of stimulus modality and learning on the cognitive processes that are involved in temporal generalization. We hope this advance in the behavioral analysis of temporal generalization will facilitate future exploration of this task, both at the behavioral level and as a basis for relating neural activity to the underlying cognitive processes, similarly to what we have shown in temporal bisection (Ofir & Landau, 2022).

## Acknowledgements

The authors would like to thank Michal Shoham and Shir Nehamkin for assistance in the design of the experiments. We thank Michal Shoham, Shir Nehamkin and Vanessa Kibel for assistance in data acquisition. We thank the members of the Brain Attention and Time Lab for their input on the work. The Brain Attention and Time Lab (PI: A.N.L.) is supported by the James McDonnell Scholar Award in Understanding Human Cognition, ISF grant 958/16. This project has received funding from the European Research Council (ERC) under the European Union’s Horizon 2020 research and innovation programme (grant agreement no. 852387).

## Supplemental Data

**Figure S1:**
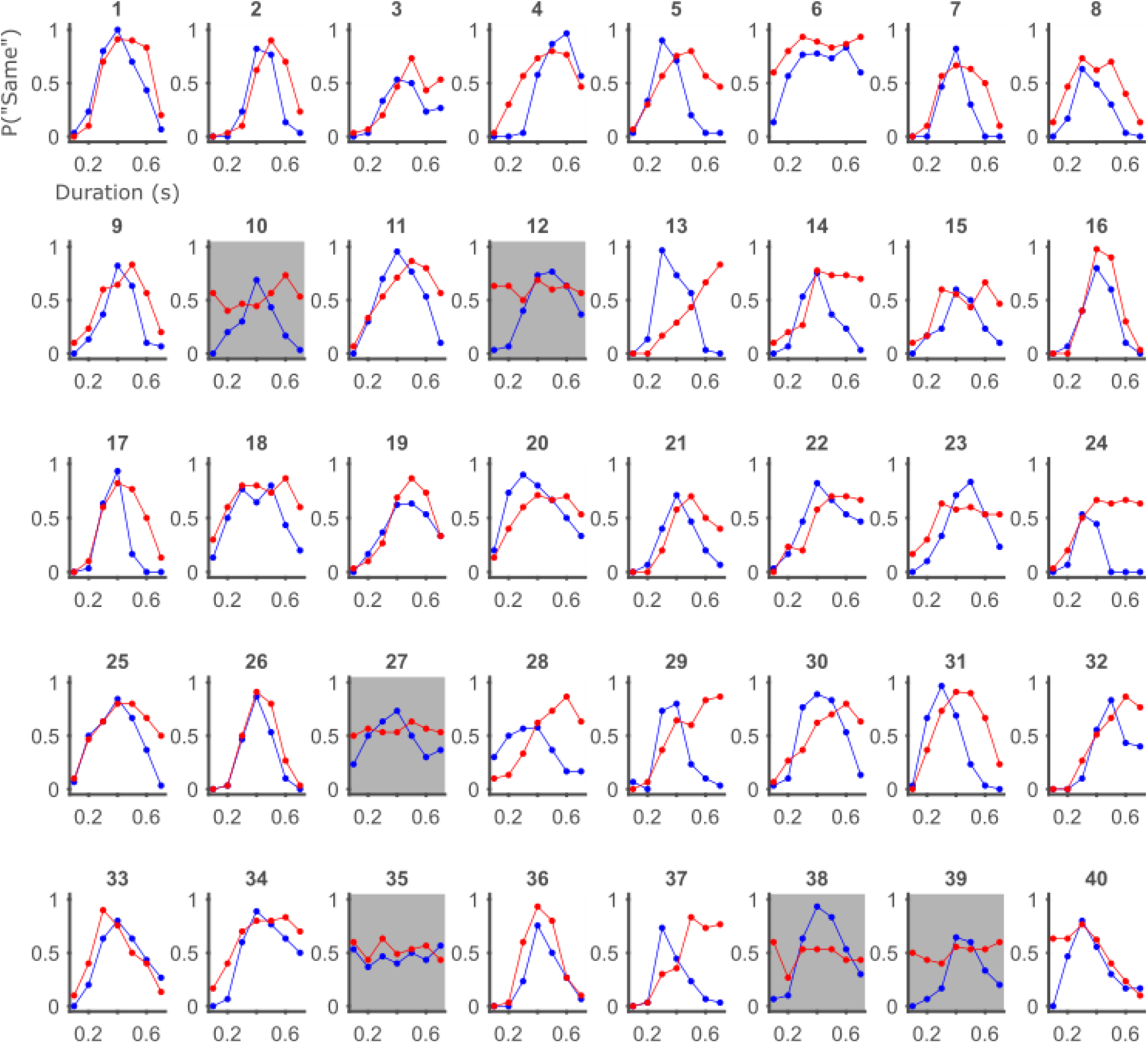
Single participants data, auditory-visual experiment. Color marks the modality (blue for auditory and red for visual). Participants with gray background were excluded from the analysis.

**Figure S2:**
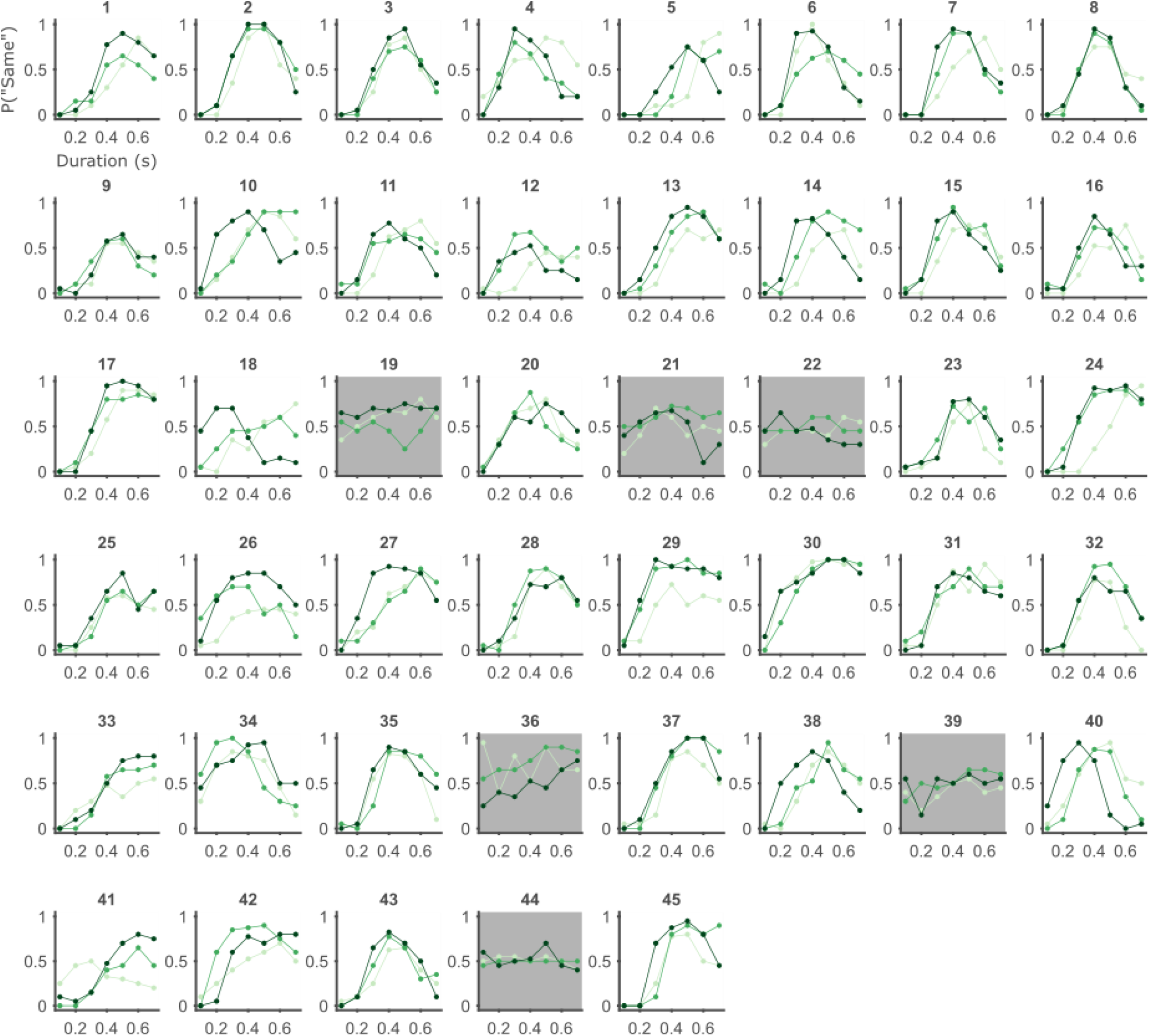
Single participants data, learning experiment. Color hue marks the tertile (light for early in the experiment and dark for late). Participants with gray background were excluded from the analysis.

## Appendices

### Appendix 1: Psychophysical function derivation for the MCG model

The MCG model assumes the following decision rule:

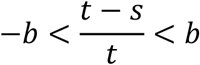

*t* is the true current presented interval. *s* is a memory sample of the standard, drawn from a Gaussian distribution with mean equal to the true standard and standard deviation that grows linearly with standard duration. *b* is a sample of the decision boundary, which is drawn from a Gaussian distribution with a given mean and standard deviation.

Since *t* is positive, and defining *w*_*i*_ = (−1)_*i*_*Bt*, the decision rule can be rewritten as:

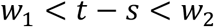

The probability of a “same” response is then

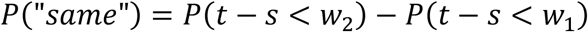

Equivalent to

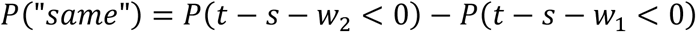

The left-hand side within each parenthesis is a sum of independent Gaussian random variables (*s* and *w*_*i*_) and a constant (*t*). Hence, that sum is itself a Gaussian random variable with the following mean and variance (*S* and *B* are the mean of memory and boundary distributions, respectively):

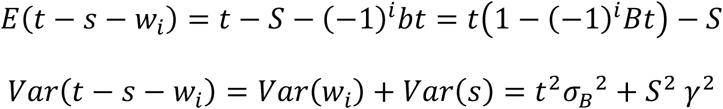

With this, we can write the psychophysical function:

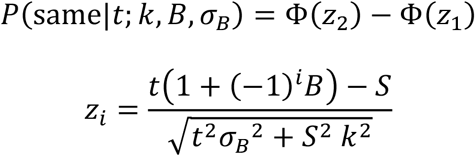

### Appendix 2: Psychophysical function derivation for the double-boundary DDM

The mathematical formulation of our model is as follows (adapted from Simen et al., 2011): A drift-diffusion process, representing the momentary perceived duration (accumulated temporal evidence), starts at the stimulus onset. The dynamics of the DV is given by:

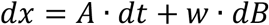

Where *x* is the DV, *A* is the drift rate (how fast the DV grows over time on average), *w* · *dB* represents the variability in momentary DV (the noise level). The probability density function of DV values at a given moment *t* is given by the following Gaussian distribution:

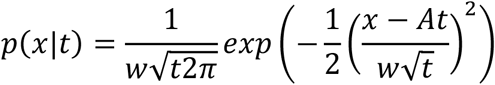

We define two additional parameters, *z*_*l*_ and *z*_*u*_, which represent the lower and upper decision boundaries, respectively. The probability of a “same” response for an in interval of duration *t* is the probability that the DV at moment *t* is between the boundaries:

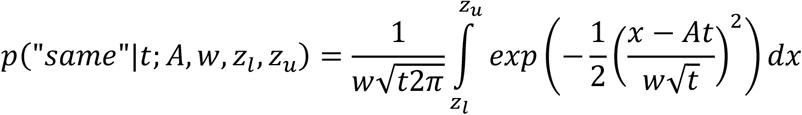

We perform the following change of variables, and define Φ(*x*) as the cumulative standardized Gaussian function evaluated at *x*:

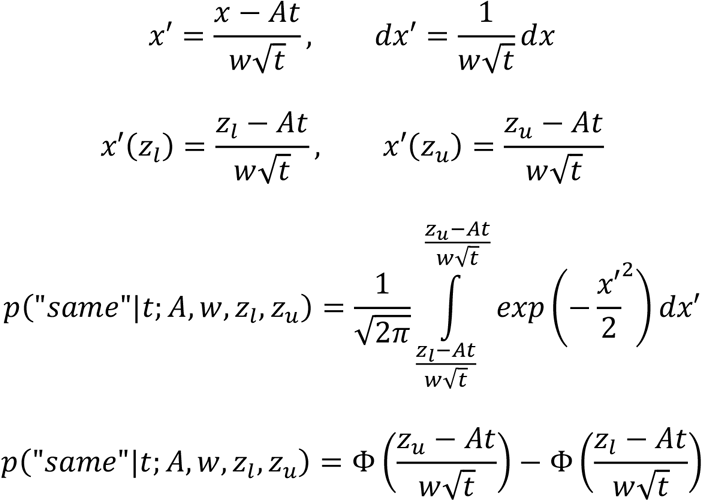

In our data it is not possible to estimate all four parameters since the boundaries and drift coefficient can always offset the effect of changing the drift rate. Therefore, we define three new parameters which are ratios of the diffusion and boundaries against the drift rate: 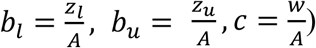 which brings us to the final psychophysical function:

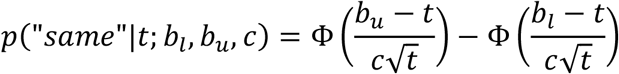

### Open Practice Statement

The data and code for all experiments and analyses are available at https://osf.io/87zbp/. Neither of the experiments were preregistered.

